# Spike Train Scalograms (STS): a Deep Learning Classification Pipeline for Neuronal Cell Types*

**DOI:** 10.1101/2025.02.10.637424

**Authors:** Gianluca Amprimo, Lorenzo Martini, Begum Bilir, Roberta Bardini, Alessandro Savino, Gabriella Olmo, Stefano Di Carlo

## Abstract

Classifying neuronal cell types is crucial for understanding the intricate circuitry of the cerebral cortex, which comprises diverse specialized neurons essential for brain function. Traditional Machine Learning (ML) approaches rely on manually engineered electrophysiological (EP) features, often overlooking subtle and complex patterns within spike train data. This study introduces a novel Spike Train Scalograms (STS)-based Deep Learning (DL) pipeline that integrates Continuous Wavelet Transform (CWT) scalograms with pre-trained Convolutional Neural Network (CNN) architectures to classify neuronal cell types with high accuracy. Utilizing patch-clamp EP recordings from 5,590 murine cortical neurons, the pipeline transforms spike trains into time-frequency representations via CWT, capturing both transient and sustained signal characteristics. These scalograms are then processed by fine-tuned CNN architectures, including *InceptionV3*, which achieved a balanced accuracy and weighted *F1*-Score of 90.53% and 90.03%, respectively. The STS pipeline effectively distinguishes between major neuronal types such as Pvalb, Sst, Vip/Lamp5, and Excitatory neurons, even in the presence of class imbalances. Moreover, an explainability analysis using saliency maps and SHAP revealed high correspondence between the DL approach, the ML baseline and biological knowledge of these neuronal types. The results demonstrate that, by combining an advanced spectral analysis with DL techniques, neurons can be classified with high accuracy, employing only two raw sweeps rather then the full stimulation range required by shallow approaches.

## I. INTRODUCTION

The cerebral cortex is an intricate biological system characterized by a remarkable diversity of specialized cells that form the neuronal circuits fundamental for brain functions [1]. Brain circuits’ fundamental neuronal elements consist of cells out of two major types, distinguished by the neurotransmitter used in their neural connections: Glutamatergic (Excitatory) and GABAergic (Inhibitory) neurons [2]. The balance between those two classes guarantees the correct function of highly complex neural processes. In addition, such classes subdivide into many subtypes, differing in morphology, transcriptional profiles, and electrophysiological (EP) properties and behavior [3] [4].

Electrophysiology can elucidate how neurons respond to and send stimuli, a crucial aspect of inter-neuron communication and network functionality. Researchers can record ion channel activity and neuronal EP state in real-time under controlled conditions by employing the patch-clamp technique [5] to investigate these aspects. When stimulated, a neuron can generate an Action Potential (AP) [6], which propagates to connected neurons, transmitting information through the network. Neural information is encoded by single APs and by the rapid sequences of APs, known as spike trains, enabling complex signaling. Slight variations in neuronal responses are vital for differentiating neuron subtypes [7] and understanding their specific roles [8], underscoring the importance of accurate classification approaches. Patch clamp experiments usually require tens of different stimulations to obtain a complete characterization of a single neuron, increasing the complexity of the experimental setup. Moreover, traditional data processing methods exploit only in part the rich information hidden within neuronal EP traces, by defining knowledge-based, human-engineered features. These features are indeed helpful for a first-level characterization. However, they do not allow for the appreciation of nuances and patterns in the signal that are fundamental for the dynamic balance of neural networks.

On the other hand, recent advances in Artificial Intelligence (AI) and Machine Learning (ML) enable automated analysis and the discovery of subtle gating patterns, thereby broadening the scope of the study of electrophysiology [9]. Combining Spectral Analysis (SA) with Deep Learning (DL) overcomes manual, knowledge-based feature creation by automatically extracting hierarchical, nonlinear patterns and improving scalability and robustness.

Therefore, this work introduces the Spike Train Scalograms (STS) pipeline, which uses DL and the Continuous Wavelet Transform (CWT) to analyze EP responses of cortical neurons and classify them into different neuronal types. The strength of this STS pipeline stems from its ability to directly process the raw neuronal response, retaining relevant information that may not be captured by knowledge-based and high-level EP features. The STS pipeline employs well-known pre-trained Convolutional Neural Networks (CNNs), fine-tuned to classify the neurons into the main brain cortex neuronal types. This work considers, optimizes, and compares different DL architectures to find the best model for the STS pipeline. Since this work is the first to employ a substantial number of neurons for training, a shallow ML model has been computed to establish a baseline for the feature-based approach for comparison.

The provided STS pipeline performs as well as the conventional shallow approach and better then the literature DL models in classification accuracy, and does not require time-consuming experimental procedures to extract high-level features. An explainability study reveals that the STS pipeline can appreciate the valuable information from the full spike train signal, while remaining consistent with knowledge-based features representation.

## II. BACKGROUND

### A. Related works

Over the past decades, as outlined in [9], a variety of Machine Learning methods have been employed to classify neuronal cell types based on EP recordings. Early studies predominantly relied on knowledge-based, high-level features derived from APs and spike trains, feeding these into shallow ML models, such as k-Nearest Neighbors, Gaussian Mixture Models, or Random Forests, to differentiate broad neuronal classes and subtypes.

On this line, Gouwens et al. [3], using the Allen Cell Types Database [10], leveraged morphological and electrophysiological knowledge-based features (e.g., AP threshold, amplitude, inter-spike intervals) for clustering analysis, identifying 17 neuronal types. Their later work [4] combined unsupervised clustering of morphological and electrophysiological features with supervised classification to predict transcriptomic types. Similarly, ISICA [11] used gamma distribution shape factors and inter-spike interval variations to classify dopaminergic or CA1 pyramidal neuron subtypes. While these approaches established a foundation for systematic neuron classification, they depend on high-level feature definition, which limits their capacity to capture subtle, complex patterns in neuronal activity.

More recent investigations have explored DL techniques. Ophir et al. [12] introduced a DL framework achieving higher accuracy than shallow ML, leveraging both EP features and domain adaptation to generalize across species. Although powerful, this DL-based method relies on raw temporal signals or simple spectral representations, potentially missing essential time-frequency details.

Some works enrich feature representations by combining SA and ML. Ghaderi et al. [13] applied the Discrete Cosine Transform to *in vivo* patch-clamp recordings combined with shallow ML classifiers. Wang et al. [14] combined the Fourier transform and CNN-based DL models, incorporating frequency-domain information. However, these methods did not fully exploit time-frequency localization, as they relied on basic spectral transformations rather than more sophisticated time-frequency decompositions like the CWT (see Section II-B).

In summary, existing approaches have made significant progress. Still, the state-of-the-art lacks methods to capture complex and non-stationary neuronal signal patterns. This work integrates a robust time-frequency decomposition with DL-based feature extraction, simultaneously leveraging advanced spectral transformations and DL architectures to reduce manual intervention and produce more discriminative representations. Indeed, the STS pipeline moves in this direction, employing CWT for time-frequency representation and DL architectures capable of uncovering intricate, nonlinear relationships in neuronal signals.

### B. Continuous Wavelet Transform

The CWT is a mathematical method for analyzing signals by providing a time-frequency representation that emphasizes how frequency components evolve. The CWT employs wavelets (i.e., localized oscillatory functions with finite duration) that can be scaled and translated over time. This flexibility allows the CWT to adapt its resolution dynamically, offering better insights into non-stationary signals where frequency content changes over time [15]. The CWT is defined mathematically as:

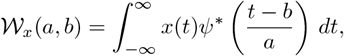

where *a >* 0 is the scale parameter, inversely proportional to frequency, *b* represents the time-shifting parameter, and *ψ*(*t*)is the mother wavelet. The mother wavelet is the prototype function scaled by *a* and shifted by *b* to create a family of wavelets adapting to different signal characteristics. Mother wavelets used in the CWT are diverse and application-specific, as their choice determines the resolution and sensitivity of the analysis to specific signal features. For example, the *Mexican hat wavelet*, derived as the second derivative of a Gaussian function, is effective for detecting singularities or abrupt transitions [16].

The result of the CWT is a two-dimensional array of coefficients 𝒲_*x*_(*a, b*), which describes the signal’s frequency content at specific times. These coefficients can be visualized using a *scalogram*, i.e., a two-dimensional representation of the energy distribution across time and scale (i.e., frequency).

CWT transforms a one-dimensional signal into a 2D matrix form. For this reason, it combines well with CNNs for solving classification or regression tasks, and this approach already proved effective over non-stationary biological signals [17]. For instance, Ruffinatti et al. [18] employed CWT to investigate spatial patterns in calcium oscillations within developing neurons. Although this study focused on the spatial compartmentalization of calcium signals rather than EP classification, it demonstrates the power of CWT in capturing complex neuronal dynamics. This rationale motivated the usage of CWT in the STS pipeline, as described in Section

### C.Transfer Learning

Most patch-clamp datasets provide dozens or hundreds of cells [19], and even the largest-scale EP experiments may not produce more than a few thousand labeled cells [3], [4]. Given this limitation in EP data availability, *transfer learning* is central to developing DL models for cell type recognition. Transfer learning is a technique in DL that applies pretrained models to new tasks by leveraging knowledge learned on larger datasets [20]. This approach reduces the need for extensive computational resources and large labeled datasets, primarily when the new task is related to the original training task. This is possible thanks to *fine-tuning*, i.e., adapting the pre-trained model by updating its parameters through the limited labeled data available from the new task. This procedure involves maintaining the *feature extraction* part of the network (e.g., the convolutional layer of a CNN). In contrast, the final classification layer is replaced with one suitable for the target task, matching the number of output classes. This approach is broadly applied in different application domains, such as medical imaging [21].

## III. MATERIALS AND METHODS

### A. Datasets

This work employs the combination of two datasets, both available in the *Allen Brain Atlas Cell Types Database* [10]: a patch-clamp dataset [3] including EP data from 1,938 neurons in the visual cortex of adult mice, along with morphological reconstructions for 461 of these neurons; a patchseq dataset [4] which combines patch-clamp with single-cell RNA sequencing (scRNA-seq) to collect EP and transcriptomic profiles of 4,200 mouse visual cortical GABAergic interneurons. Their combination contains a mix of inhibitory and excitatory neurons. The inhibitory neurons are divided into three main subtypes: Pvalb, Sst, and Vip/Lamp5, based on the transcriptomic taxonomy [22] and the samples’ cell lines [23]. The dataset presents a strong imbalance between these classes, with excitatory neurons being a substantial minority. After data cleaning, 5590 neurons with suitable recordings and clear identities are retained.

### B. STS Pipeline

The STS classification pipeline is visualized as a sequence of processing blocks in Figure 1. The pipeline leverages the scalograms obtained through the CWT and the fine-tuned deep convolutional models (see Section III-C) to classify neuronal cells according to the spike trains emitted under stimulation. The STS implementation required to reproduce the results presented in this paper is publicly available at https://github.com/smilies-polito/spike-train-scalograms.

**Fig 1.**
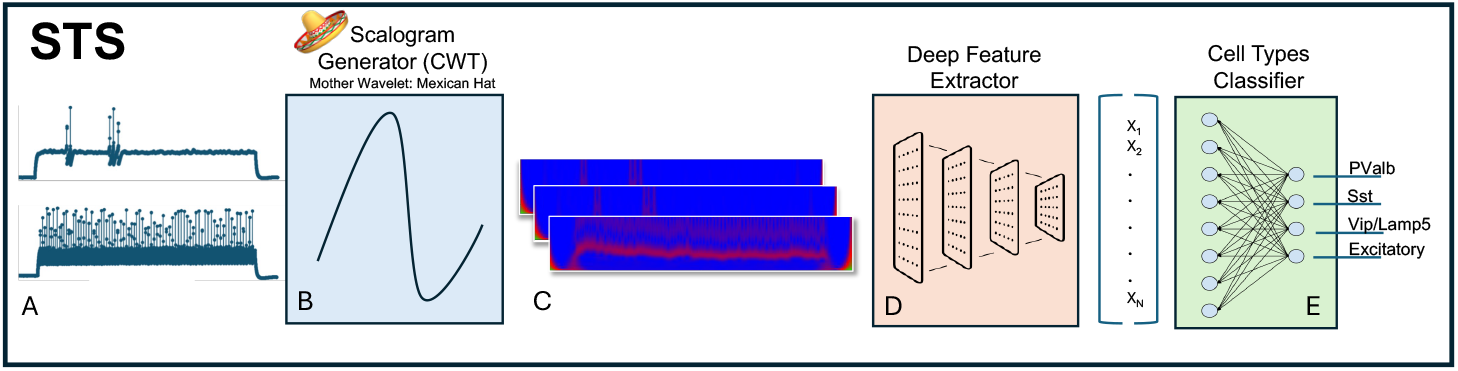
STS pipeline.

This work considers the neurons’ responses to *long square* current clamp stimulations [23]. Each stimulation recording is identified as a *sweep*. Specifically, the pipeline considers the electrophysiological response of each neuron to two specific long square sweeps:

1. the sweep corresponding to the rheobase;
2. the sweep corresponding to the rheobase +40 pA.

The *rheobase* is the minimum current amplitude of a long-duration electrical current required to initiate an AP in a specific neuron. This property reflects the cell’s inherent excitability [24]. These two sweeps are thus chosen as representative of the overall neural response to a long square current stimulation. The choice of rheobase-centered sweeps from long square stimulations [10] has solid grounds: the long square stimulation duration is sufficient to observe the instantaneous response of the neuron, as well as possibly adaptation, refractory periods, and burst events [24]; a focus on the rheobase-centered sweeps highlights the first excitable response of the neuron. The rheobase sweep covers the minimum stimulus the neuron responds to, informing of its excitability. However, the rheobase usually gives rise to a single or, at most, a few spikes. Therefore, analyzing the train behavior requires stronger stimulation from higher-amplitude current stimuli, that is, sweeps above the rheobase. Experiments do not test and record all current steps but rather a finite but inconsistent number of suprathreshold stimulations per neuron. By a quick inspection of the datasets, the sweep at rheobase +40 pA is the most available supratreshold stimulation, allowing the highest number of neurons to be retained. Notable differences can be observed in the steepness, shapes, and distribution of spikes inside the response to these two sweeps across the samples. For instance, Figure 2 shows the burst behavior followed by an abrupt quiescent period in Pvalb inhibitory neurons (green), in contrast to the regular spiking pattern in Excitatory neurons (blue).

**Fig 2.**
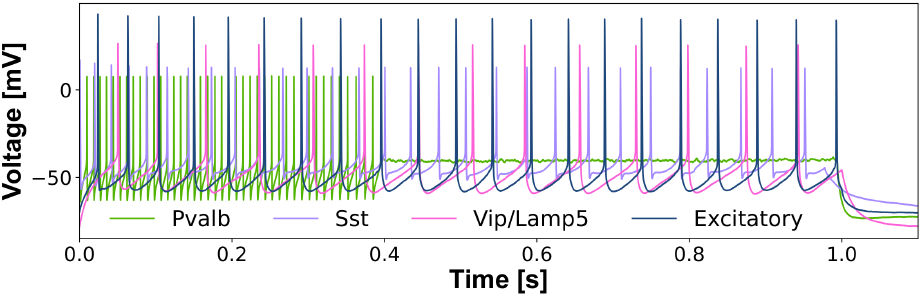
Spike trains of the four classified cell types referring to the rheobase +40 pA stimulation.

These two raw response signals to two sweeps in the time-voltage plane are the input of the STS pipeline (Fig. 1). A short description of each component follows.

#### a) Scalogram Generator

The first operation of the pipeline generates the scalograms for the selected sweeps through the CWT of the electrophysiological responses (Fig. 1A). The raw signals are downsampled as a preliminary step since voltage responses are not uniformly sampled during the patch-clamp experiment (200 - 50 kHz). Still, this parameter depends on a technical difference between years of recordings [23]. Downsampling not only uniforms data repre-sentation but also reduces computational effort for computing CWT. After carefully inspecting the spectra and considering the reconstruction error due to the downsampling, a final sampling frequency of 4 kHz was selected to compromise spectral resolution and efficiency to generate homogeneous representations of all cells.

Application of the CWT and generation of the scalogram follows (Fig. 1B). Regarding the CWT parameters, this pipeline uses the *Mexican hat* as the mother wavelet because of its capability of detecting abrupt transitions (see Section II-B), which are peculiar in the case of burst events followed by a quiescent period characteristic of some neurons. Moreover, the scalograms (Fig. 1C) are restricted to include only the frequencies between 20 Hz and 200 Hz, which retained most of the spectral power of the spike trains in the data. This smaller range also reduces computational resources required for computing CWT.

#### b) Deep Feature Extractor

A deep feature extractor based on CNNs (Fig. 1D) processes the scalograms to extract relevant information for the classification of cell types. As the samples available for training the models are limited (4472 cells), this work exploits convolutional models for image recognition, pre-trained on the ImageNet dataset. While the final classification task is different, the features that this model learned to extract from images during pre-training may also provide a solid representation for the classification of scalograms. Since image recognition models expect a three-channel input matrix, a replica of the rheobase scalogram is concatenated as the third channel for input compatibility. Moreover, scalograms are resized and transformed according to the input transformation required by the pre-trained model. This procedure overall encompasses rescaling and normalization. As the pipeline is built using <monospace>PyTorch</monospace> [25], this work compared some popular and maintained pre-trained models available in its *model-zoo* [26] as deep feature extractors, namely *MobileNetV2, ResNet18, DenseNet121* and *InceptionV3*. The processed CWTs and the trained best models for each architecture are stored on Kaggle and publicly available at https://www.kaggle.com/datasets/smiliesatpolito/STS-data.

#### C. Cell Types Classifier

The feature maps produced by the CNN architecture are squeezed (i.e., concatenated along one single dimension), generating a feature vector which is then classified by a fully connected layer (Fig. 1E) with four neurons, each predicting the probability for one of the four considered neuronal types (i.e., Pvalb, Sst, Vip/Lamp5, Excitatory). During model training, dropout was applied before the classifier to derive a more robust feature representation according to which classification is performed.

### C.Training and Optimization

This work followed a holdout strategy to train and test the STS pipeline. Therefore, after combining the two datasets described in Section III-A, the data were divided into train and test sets with an 80%-20% criterion (i.e., 4472 cells in train and 1118 cells in test), stratifying samples such that the two sets maintained the same distribution of classes. Then, the classification model was trained by exploiting a 3-fold-cross-validated Bayesian search [27] to identify the optimal hyperparameters for classifying cell types. For each proposed configuration, 3-fold cross-validation (3-fold-CV) evaluates the model by splitting the dataset into three subsets. The model is trained on two subsets and validated on the third, repeating this process three times to compute an average performance metric. This approach ensures a robust evaluation of hyperparameters’ configurations while reducing the likelihood of overfitting a single validation split. As cell type distribution was unbalanced in the training data, balanced accuracy (i.e., accuracy weighted by class numerosity [28]) was the objective function to maximize during the optimization. Several runs took place to compare the efficacy of different deep feature extractors. DL models were optimized using the Adam algorithm with a learning rate decay policy. Pre-trained weights of the deep feature extractor block were used as initialization for the network, allowing complete retraining (i.e., no frozen layers).

### D. Pipeline Evaluation

Evaluating four deep feature extractors led to comparing four optimized configurations of the STS pipeline on the holdout test set. Expressive performance metrics were computed for unbalanced multi-label classification problems, such as *F1*-score and balanced accuracy. Given the state-of-the-art proposes classification solutions based on human-engineered features, a baseline shallow ML model was trained on EP features to assess the improvement brought by combining scalograms with convolutional deep feature extractor. In particular, a tree-based ensemble classifier, namely the *Light Gradient Boosting Machine (LGBM)*, was trained using the EP features exposed by the two datasets. To ensure a fair comparison, baseline shallow ML model training included only EP features derived from responses to the long square stimulation in the suprathreshold phase (i.e., above rheobase), related to both the single spike response (i.e., voltage and time values at threshold, peak, and through of the first evoked action potential) and the inter-spike response. The latter includes features specifically connected to spike trains, such as the slope of the frequency-current (f-I) curve, average inter-spike interval (ISI), adaptation, latency, presence of burst, pause, and delays [23]. Indeed, the two scalograms selected as input to the STS pipeline should reflect these properties, plus possibly additional characteristics in the frequency domain improving the final classification. Training and optimization of the baseline model relied on the methods described in the previous section (i.e., Bayesian search with 3-fold-CV).

### E. Explainability

A final explainability analysis of the STS pipeline and the LGBM baseline revealed which deep and high-level features these models leverage during classification.

For the deep feature extractor of the STS pipeline, saliency maps [29] described what parts of the input scalograms were more significant during classification. Saliency maps are generated by calculating the output gradient with respect to the input image pixels. The magnitude of these gradients indicates which pixels most influence the prediction. By visualizing these values, saliency maps highlight the key regions that impact the classification. This work provides saliency maps for four cells in the test set, representing the neuronal types under investigation.

For the shallow baseline, explainability analysis established the EP features that drove the prediction of each class through SHAP [30]. This approach, based on *game theory* (i.e. Shapley values), assigns feature importance by calculating the average contribution of each feature to the prediction.

## IV. RESULTS AND DISCUSSION

### A. Classification performance

Figure 3 compares the mean training and validation losses for the four optimal configurations of the STS pipeline according to Bayesian optimization. Validation and train loss follow a decreasing trend for all configurations, denoting progressive learning of the cell types classifier. As it can be appreciated, a reasonable and often negligible gap exists between paired training and validation losses. All variants appear to plateau after 15 epochs, denoting a fast convergence likely thanks to an effective knowledge transfer from the pre-trained models.

**Fig 3.**
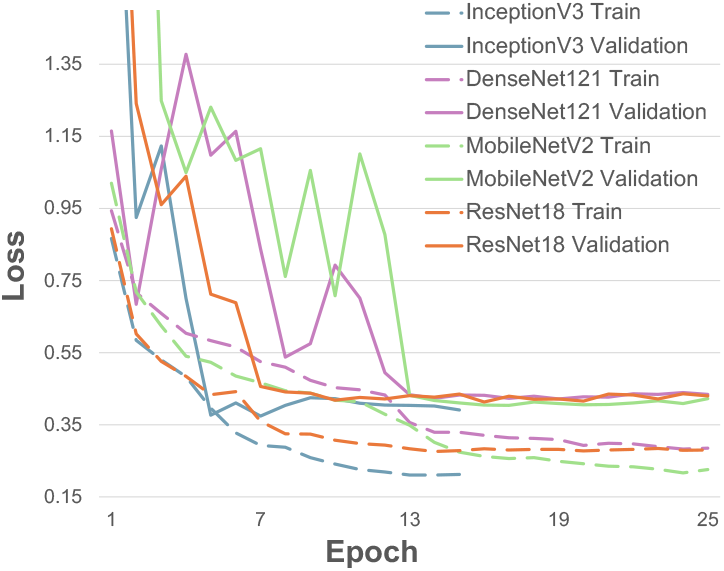
Mean losses over 3-fold-CV of the optimized STS pipelines, employing different pre-trained deep feature extractors. Dotted notation is used for training losses, whereas bold lines refer to validation losses.

Figure 4 reports the bar plots visualizing the mean and standard deviation of balanced test accuracy. This metric was maximized during the 3-fold-CV Bayesian search for the four trained configurations plus the baseline shallow ML model. Side-by-side, Figure 4 reports the results on the holdout test. Overall, all the configurations of the pipeline achieve very high and comparable performance (above 88%) in both the cross-validation and the holdout dataset, coherently with the trends observed in the train and validation losses. Holdout test values fall within the confidence range of crossvalidation. Thus, the cell type classifiers avoided overfitting.

**Fig 4.**
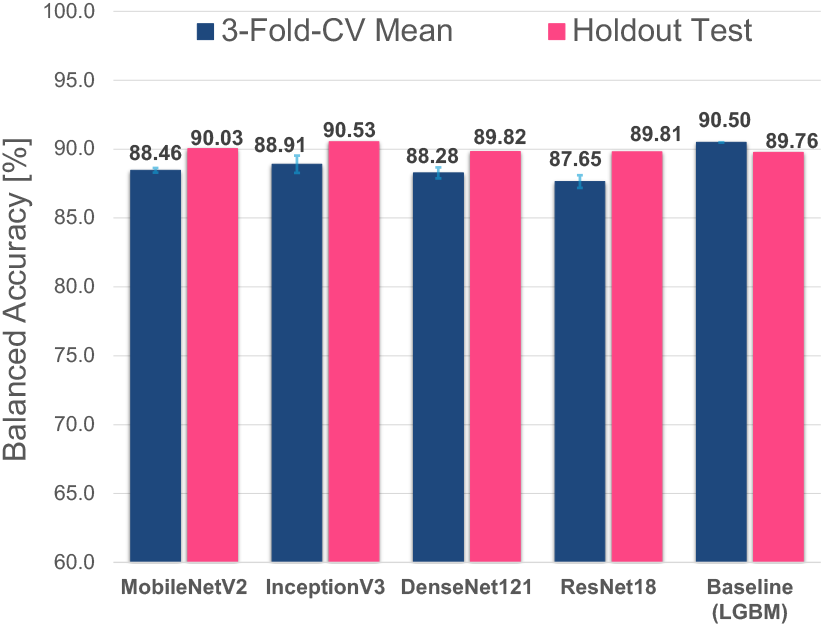
Balanced accuracy of optimized configurations of the STS pipeline during 3-fold-CV Bayesian optimization (mean value as a blue bar, plus standard deviation as an error bar on top) and during classification on the holdout test data (pink bar).

In addition to balanced accuracy, the F1-Score is analyzed. All the configurations scored weighted F1-Score above 88% on the holdout test, in particular: 89.72% *MobileNetV2*, 90.03% *InceptionV3*, 89.27% *DenseNet121* and 89.81% *ResNet18*. Again, values were comparable to the shallow baseline, which had 89.00% F1-Score.

These results imply that all the DL architecture considered as deep feature extractor learned a similar and effective data representation, so are basically interchangeable in the STS pipeline. Among them, *InceptionV3* appears to slightly surpass the others, as well as the shallow baseline.

Comparing the STS pipeline with DL-based classification models in the state of the art, the most similar works are those from Ophir et al. [12] and Wang et al. [14]. However, they both performed training and validation on a sub-portion of the data employed in this work (the patch-clamp dataset from [3]), thus less than 2,000 neurons, considering both training and testing. Instead, the results of this work were achieved on a total of 5,590 cells, of which 1,118 were kept as a completely unseen holdout test set. Considering this aspect and the different approaches used for model validation (i.e., cross-validation versus holdout testing), the comparison of the results should be interpreted cautiously. Despite these observations, the STS pipeline using *InceptionV3* surpasses the one from Ophir et al. [12], with an additional 3% in weighted F1-Score (87.7% vs 90.53%). Regarding the work from Wang et al. [14], which also investigated the use of spectral properties for classification but limited to the first AP, also in this case, the STS pipeline had superior performance (*F1-Score*:88.75% reported by [14]).

Compared to the shallow LGBM model trained on all the cells available in this study, the CWT-based approach of the STS pipeline does not cause any loss in classification accuracy. Therefore, the data representation learned using CWT and CNN is superior to the other DL models in the literature and equivalent to the established, high-level EP features.

However, the scalogram-based representation provides two main benefits compared to high-level feature. On the one hand, it reduces experimental resources required for the classification, as only two sweeps must be recorded, including the rheobase. On the other hand, it provides a more in depth description of neuronal activity, as further supported by the explainability analysis.

### B. Explainability analysis

The explainability analysis focused on the pipeline embedding *InceptionV3* to compare the representation learned by the best STS pipeline against that of the optimal baseline model. Figure 5 visualizes simultaneously the saliency maps of two representative cells from *Excitatory* and *Pvalb* types, along the per-class summary plots provided by SHAP for the shallow classifier. Positive SHAP values (x-axis) imply that a low or high feature value predicted the class. In contrast, a negative value drove the classification towards one of the other classes. Similar plots and observations for *Vip/Lamp5* and *Sst* cells are available at https://github.com/smilies-polito/spike-train-scalograms but are omitted here for brevity. The saliency maps show that for *Excitatory* cells, the model concentrates on the initial signal portion, specifically around the first spike and at low frequencies. This suggests that the first spike is crucial in driving classification for this cell type, in line with the steady firing pattern typical of *Excitatory* neurons. In contrast, for *Pvalb* cells, the saliency is distributed across a broader portion of the signal, highlighting upper regions of the CWT corresponding to high-frequency components. Notably, the model emphasizes the regions where the spike train burst ends and starts the quiescent period, which aligns well with the fast-spiking behavior typical of *Pvalb* neurons. This differential attribution suggests that the classifier captures relevant dynamical features specific to each neuronal subtype, reinforcing the biological significance of the learned representations. Moreover, both cases are coherent with the top five most important features highlighted by the SHAP analysis of the shallow model. For *Excitatory*, the features *fast trough t, fast trough v* and *trough t* characterize the shape of the first spike, coherent with what highlighted by the saliency. In contrast, for *Pvalb*, the *f i curve slope* and *adaptation* characterize the spike train dynamic and its abrupt changes thus weighting the whole spike train behavior.

**Fig 5.**
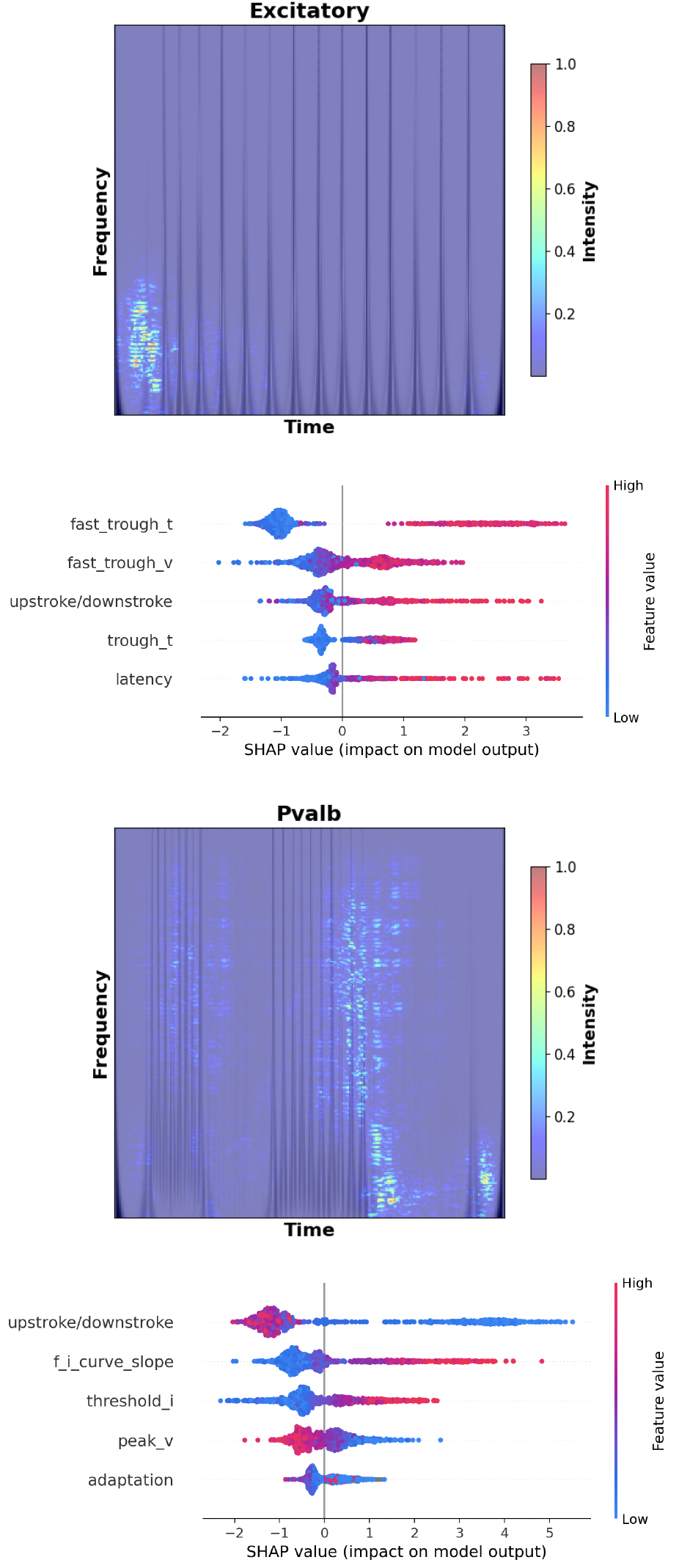
Interpretability of STS pipeline using saliency maps versus interpretability of shallow baseline using SHAP.

While converging on the biological significance, the CWT scalograms provide a more in-depth visualization of regions in the time-frequency neuronal response that are relevant to the cell types dynamics, with respect to the broad and shallow description provided by EP features.

## V. CONCLUSIONS

This study introduces STS, a robust pipeline that integrates CWT scalograms with fine-tuned DL architectures for accurate neuronal cell type classification. By capturing intricate time-frequency patterns in spike trains, the pipeline retain relevant information to classify neuronal types and provides a more expressive data representation compared to shallow ML models using high-level EP features. Moreover, only two experimental sweeps are needed, whereas EP are the results of averaging several sweeps above rheobase.

It is relevant to highlight how the classification distinguishes subtypes, usually characterized at the transcriptomic level, from their EP profiles alone. Indeed, EP experiments usually require secondary experimental analysis, such as creating transgenic mice lines supporting the pre-recording identification of specific neuron populations or transcriptional profiling of the neurons. Both cases involve expensive technologies and procedures, reducing the feasibility of such studies. A purely EP classification would allow, in the first instance, a first-level ground truth for deeper investigations without increasing the experimental expenses.

This work is a first step towards building a more comprehensive computational framework providing complementary solutions for analyzing, characterizing, and modeling brain heterogeneity based on multi-modal neuronal data. In this direction, an interesting future endeavor includes testing the STS pipeline on human neurons to address possible issues in domain generalization across species, as done by [12]. Overall, the STS pipeline demonstrates the power of DL-based solutions in neuroscience, offering a scalable and automated method to harness the complex dynamics of neuronal electrophysiological responses, accurately capturing the heterogeneity of neuronal cell types.

## ACKNOWLEDGEMENTS

The authors acknowledge the use of GPT-4.0 and GPT-4.0 with optimization for language and text refinement during manuscript preparation. The scientific content, data analysis, and interpretations remain the authors’ sole responsibility.

